# Optical clearing of living brains with MAGICAL to extend *in vivo* imaging

**DOI:** 10.1101/507426

**Authors:** Kouichirou Iijima, Takuto Oshima, Ryosuke Kawakami, Tomomi Nemoto

## Abstract

To understand brain functions, it is important to observe directly how multiple neural circuits are performing in living brains. However, due to tissue opaqueness, observable depth and spatiotemporal resolution are severely degraded *in vivo*. Here, we propose an optical brain clearing method for *in vivo* fluorescence microscopy, termed MAGICAL (Magical Additive Glycerol Improves Clear Alive Luminance). MAGICAL enabled two-photon microscopy to capture vivid images with fast speed, at cortical layer V and hippocampal CA1 *in vivo*. Moreover, MAGICAL promoted conventional confocal microscopy to visualize finer neuronal structures including synaptic boutons and spines in unprecedented deep regions, without intensive illumination leading to phototoxic effects. Fluorescence Emission Spectrum Transmissive Analysis (FESTA) showed that MAGICAL improved *in vivo* transmittance of shorter wavelength light, which is vulnerable to optical scattering thus unsuited for *in vivo* microscopy. These results suggest that MAGICAL would transparentize living brains via scattering reduction.

## Main Text

How do neurons work together for brain functions in living animals? In the brain, numerous neurons connect to each other three-dimensionally (3D) to form neural circuits underlying brain functions. Functional characteristics of neural circuits depend on neuronal activity and synaptic connectivity, both of which are reflected in fine sculptural distinctions in many cases^1-3^. Multiple neural circuits are associated with each other as a neural network, and are executed for rapid processing simultaneously, to respond to various situations and their transitions^4^. Therefore, to understand brain functions, *in vivo* imaging techniques are required to observe intact neural circuits extended into deep regions, at high spatiotemporal resolution.

Confocal microscopy and two-photon microscopy can provide 3D and time-lapse images by sequential optical sectioning in thick specimens without mechanical destruction^5,6^. However, it is difficult for the fluorescence microscopy, especially to confocal microscopy, to achieve sufficiently deep, fine, and fast imaging *in vivo*, because tissue opaqueness disturbs and attenuates both excitation lights and fluorescence signals. A simple but practical technique to overcome the opaqueness is high-intensity illumination by using high-power excitation laser, but it has a risk of causing invasive phototoxic problems via reactive oxygen species production. Thus, new strategies are needed to overcome tissue opaqueness for *in vivo* imaging.

Recently, many optical clearing methods were reported for fluorescence microscopy in fixed organs including brains^7-13^. These clearing methods enable confocal microscopy to achieve fine and deeper imaging, and have brought a paradigm shift in connectome analysis, to create connection maps over comprehensive neural networks. However, these clearing methods need non-physiological procedures to transparentize fixed organs, thus are especially inapplicable to living brains for *in vivo* fluorescence microscopy. Here, we propose a new strategy for brain-clearing in *in vivo* microscopy. Our clearing method is simply based on glycerol administration via drinking water, so we named it MAGICAL (Magical Additive Glycerol Improves Clear Alive Luminance).

## Results

### Enhanced fluorescence signals *in vivo*

For *in vivo* brain imaging with fluorescence microscopy, the conventional method involves open-skull surgery. It consists of making a cranial window, in which the light-scattering skull bone is replaced by a clear cover glass, and is conceived as a key step affecting the quality of images in day-to-day experiments^14^. The cranial window, and probably the brain itself, lose transparency easily, which may be caused by unsuitable surgical techniques, leading to bleeding and damage inside/outside the brain, and by unknown factors related with the recovery process. Thus, a smaller cranial window is biologically preferred to avoid injuring blood vessels and to minimize contact between the brain surface and the surgical materials^14^. However, a larger cranial window is optically preferred to collect even light emitted from a focal point at wide radiation angles, by using a high-numerical-aperture (NA) lens which governs spatial resolution and fluorescence intensity in imaging. Therefore, we tried to improve *in vivo* microscopy by reducing the detrimental effects of open-skull surgery. We focused on glycerol oral administration, which is known to suppress cerebral edema in neurosurgical patients and suggested to supply energy to ischemic neurons in cerebral stroke^15-22^.

First, we examined whether oral administration of glycerol improves cerebral fluorescence images of *in vivo* two-photon microscopy. To visualize neural circuits, we used adult H-line mice (Thy1-EYFP-H), which express Enhanced Yellow Fluorescent Protein (EYFP) in some of the pyramidal neurons. To avoid selection bias under uneven EYFP expression, the imaging area was selected according to the xy position where the hippocampus was observable at the most shallow depth. We captured 3D stacks as z-series of xy-images, using a two-photon microscope with 960-nm Ti:Sapphire laser light. The xy-images were acquired as 512 × 512 pixels at 1 frame per sec (fps), 2.2 μsec per pixel, which is the maximum speed in this condition.

The images of the cortical layer V (CxLV) neurons were captured with 3-μm z-step size (**Supplementary Movie 1**) and were reconstructed with depth color-coded maximum intensity projection (DccMIP). In mice without glycerol administration (control), fluorescence images showed the pyramidal neuron cell bodies with basal dendrites, but the density was poor as a consequence of weak capture conditions (**Fig. 1a**, **Supplementary Movie 1**). In contrast, in mice with 5 % (w/v) glycerol administration (MAGICAL), bright fluorescence images showed many cell bodies and basal dendrites with high density (**Fig. 1a**, **Supplementary Movie 1**).

**Fig 1.**
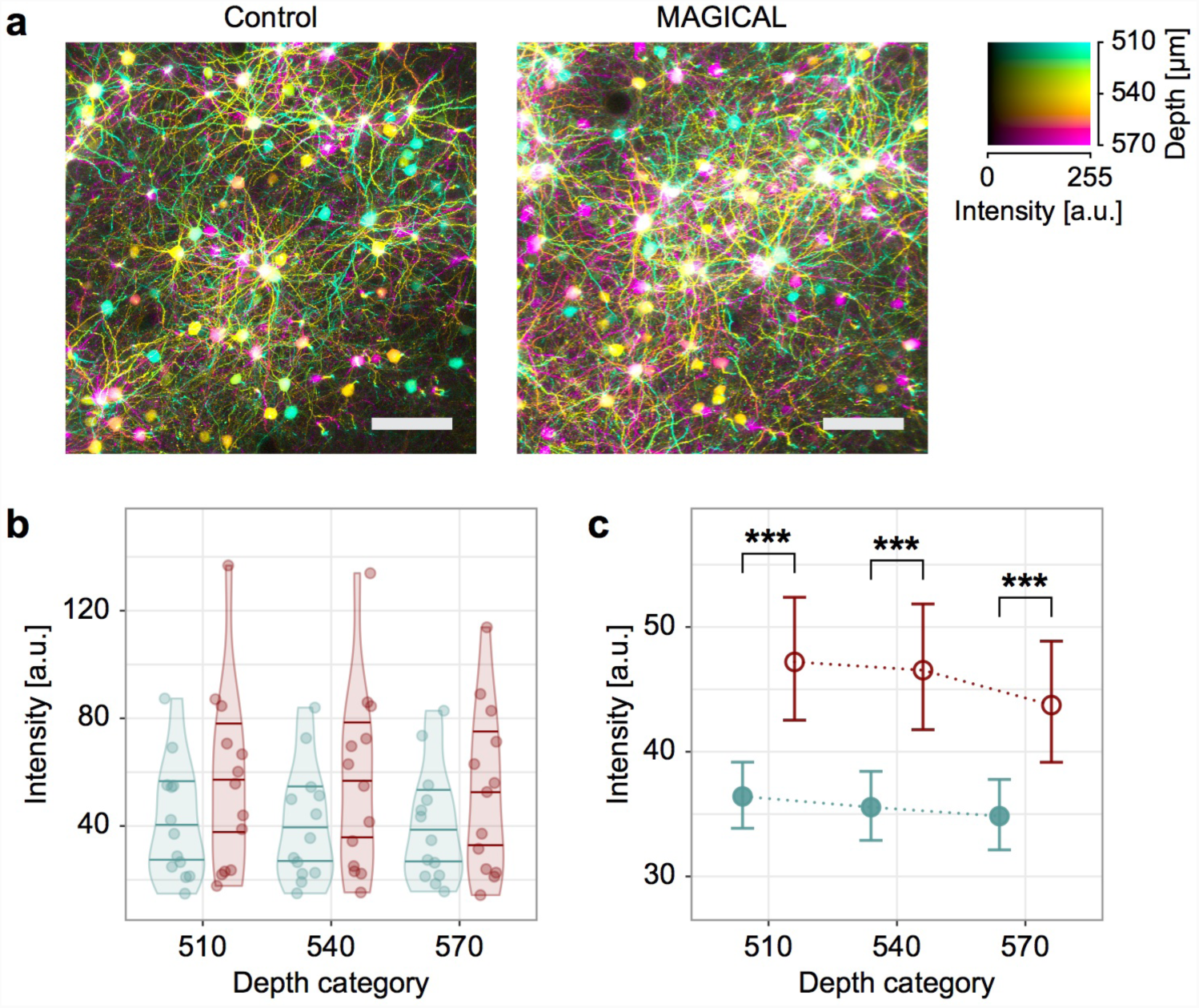
*In vivo* two-photon imaging at CxLV. (**a**) DccMIP images of CxLV at 540 ± 30 μm depth in control and MAGICAL. scale bar 100 μm. (**b–c**) Representative fluorescence intensity, predicted by the GLMM for each image stack (**b**) and “Treatment” group (**c**), along “Depth.” Distribution of plots in (**b**) is visualized with violin plots estimated by Gaussian kernel and with 0.25, 0.50, and 0.75 quantile lines. Error bars in (**c**) are asymptotic 95 % CI. Red, MAGICAL; Blue, control; *** *p* < 0.001; a.u., arbitrary unit.

In CxLV images at three different depths (**Supplementary Fig. 1**), fluorescence intensity distribution (FID) indicated that signals tended to increase with MAGICAL (**Supplementary Fig. 2**). However, it was not simple to summarize the FIDs for statistical analysis. The FIDs showed large individual differences within each treatment group, obscuring differences between the groups. Moreover, the FIDs had non-normal distributions extending beyond the detector’s dynamic range. To overcome these problems, we evaluated fluorescence intensities of pixels (FIPs) in the image stacks, by using a Generalized Linear Mixed Model (GLMM). The FIPs were randomly selected from the stack and collected as samples, if not saturation or zero intensity, and were used as a response variable in the GLMM with a Gamma error distribution and a log link function. In the GLMM, “Treatment” (MAGICAL or control) and “Depth” (3 levels) were assigned as fixed effects, and “Mouse” was assigned as a random effect for intercepts and “Depth” slopes. The GLMM comprised 929,331 FIPs in 78 image stacks, extracted from 3 depths of 26 mice under 2 treatments (*R*^2^_*GLMM(m)*_: 0.0147, *R*^2^_*GLMM(c)*_: 0.200, **Supplementary Table 1a**). In estimation for fixed effects, the GLMM indicated that MAGICAL enhanced FIPs in the CxLV images (**Fig. 1b–c**; *p* < 0.001 at each “Depth” level by pairwise comparison). These signal enhancements mean that MAGICAL can overcome tissue opaqueness enough to improve *in vivo* imaging.

### *In vivo* two-photon deep imaging with fast speed

To achieve *in vivo* imaging in deeper regions, the most effective strategy is to use much longer-wavelength, less scattered, near-infrared light at high power for two-photon excitation^23,24^. Recently, hippocampal granule cells in the dentate gyrus were visualized at 1.5-mm depth in adult mice by using a 1064 nm laser-diode light, in the most successful surgery without bleeding^24^. Furthermore, hippocampal CA1 (HpCA1) neurons can be visualized at 1.0-mm depth in adult mice, by using 1000 nm Ti:Sapphire laser light^24^. However, despite the advantage of longer-wavelength strategy, fluorescence acquisitions must collect shorter visible light, attenuated more easily by scattering and absorption in tissues depending on return distance. Indeed, both deep imaging techniques require scanning neurons at 1/16 or less fps, which is extremely slow, to visualize neural activity and to overcome image distortions caused by heartbeats. So, as a touchstone of *in vivo* deep imaging, we examined whether MAGICAL improves adult HpCA1 images at fast scanning.

By using 960 nm Ti:Sapphire laser light, HpCA1 images approximately at 1.0-mm depth were captured as 3D stacks, having xy-images (512 × 512 pixels) with 1 fps fast speed. In control mice, fluorescence images showed some dim cell bodies of pyramidal neurons (**Fig. 2a**, **Supplementary Fig. 3**). In contrast, in MAGICAL mice, fluorescence images showed many cell bodies of pyramidal neurons, some of which were clearly visualized with their partial process of apical dendrite (**Fig. 2a**, **Supplementary Fig. 3**).

**Fig 2.**
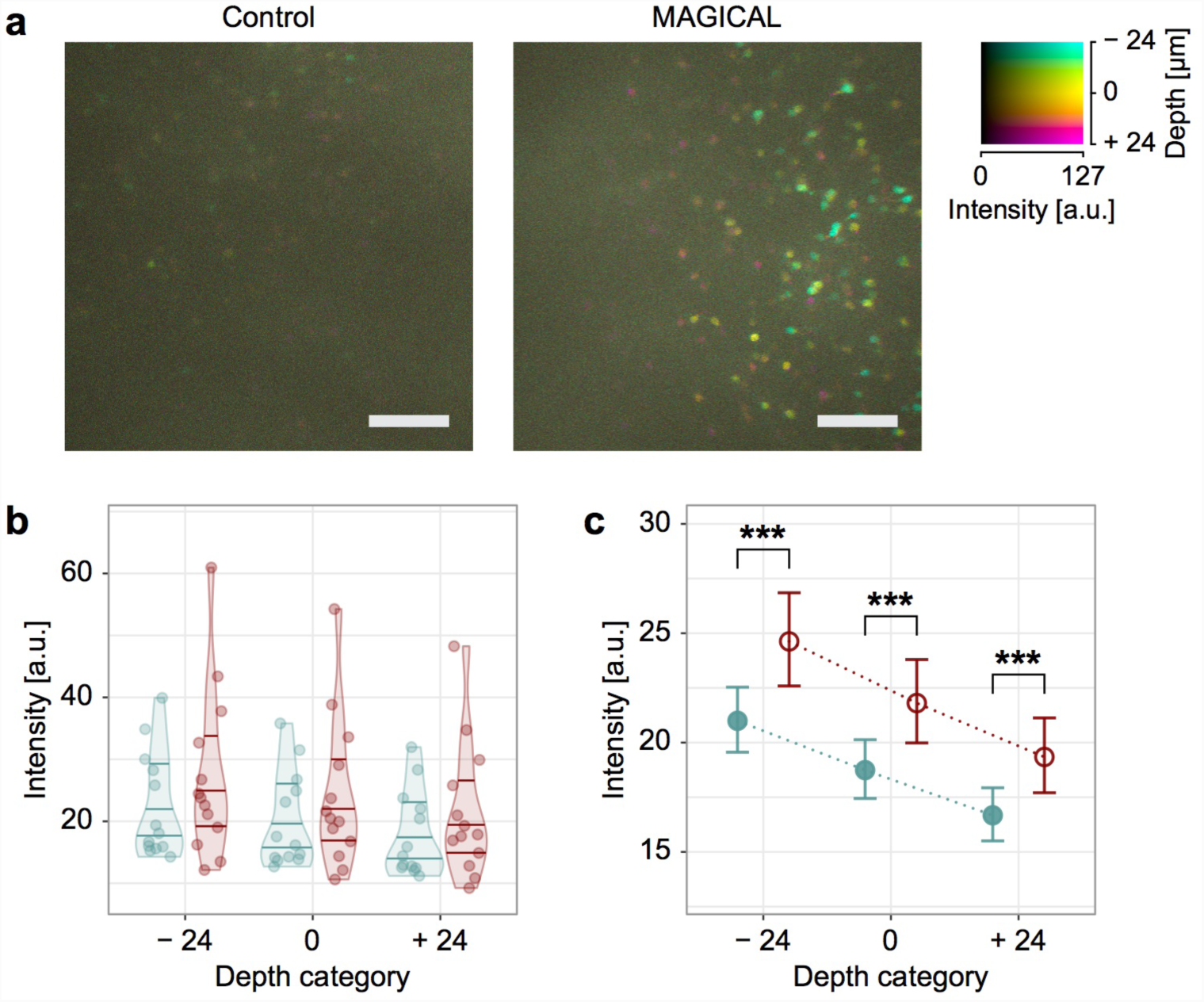
*In vivo* two-photon imaging at HpCA1. (**a**) DccMIP images of HpCA1 approximately at 975 ± 24 μm depth in control and MAGICAL. In the color code, each center depth is shown as 0 μm relative depth. scale bar 100 μm. (**b–c**) Representative fluorescence intensity, predicted by the GLMM for each image stack (**b**) and “Treatment” group (**c**), along “Depth.” Distribution of plots in (**b**) is visualized with violin plots estimated by Gaussian kernel and with 0.25, 0.50, and 0.75 quantile lines. Error bars in (**c**) are asymptotic 95 % CI. Red, MAGICAL; Blue, control; *** *p* < 0.001; a.u., arbitrary unit.

In HpCA1 images captured from three different depths ± 3 μm (**Supplementary Fig. 4**), FIDs were shifted to bright side with MAGICAL (**Supplementary Fig. 5**). GLMM at HpCA1 was comprised with 957,264 FIPs in 78 image stacks extracted 3 depths from 26 mice under 2 treatments (*R*^2^_*GLMM(m)*_: 0.0714, *R*^2^_*GLMM(c)*_: 0.190, **Supplementary Table 1b**). In estimation for fixed effects, the GLMM indicated that MAGICAL also enhanced FIPs in the HpCA1 images, despite long traversing from deep regions (**Fig. 2b–c**; *p* < 0.001 at each “Depth” level by pairwise comparison). These results suggest that MAGICAL would promote deep imaging as a practical method to visualize fast activity on mature neural circuits in adult mice.

### *In vivo* confocal fine imaging with single-photon excitation at deeper regions

Despite its superior spatial resolution, confocal microscopy is less suitable for *in vivo* imaging than two-photon microscopy, because its shorter-wavelength excitation has an inferior penetration depth and a higher risk for phototoxic effects. Moreover, confocal microscopy cannot efficiently collect fluorescence signals due to a confocal aperture, which excludes most of the signals scattered by the tissue but is needed to remove extrafocal light causing image blur. Therefore, confocal microscopy is vulnerable to tissue opaqueness, especially optical scattering. In general, *in vivo* confocal microscopy is hardly able to visualize fine neural fibers below 50-μm depth, which poses a dilemma between clear visualization and phototoxicity reduction^25^. The abovementioned effects of MAGICAL on *in vivo* two-photon imaging, suggest that it may overcome that dilemma.

Thus, we examined to what extent MAGICAL enables *in vivo* confocal microscopy to visualize neural fibers at deeper regions. We captured 3D stack images of cortical layer I (CxLI) at 50 ± 5, 100 ± 5, and 150 ± 5 μm depth in H-line mice, by using a confocal microscope with 488-nm laser light (CM image) and a two-photon microscope with 960-nm (2PM image as a reference). The 3D stack images comprise 1-μm-step z-series of xy-images, acquired as 512 × 512 pixels at 1 fps. In control mice, dim CM images at 100 and 150 μm depth were hardly able to visualize fine neural fibers except for thick shafts of apical dendrites which extend along the z-axis direction (**Fig. 3a**, **Supplementary Fig. 6**). However, in MAGICAL mice, bright CM images even at 100 and 150 μm depth showed a lot of thin neuronal processes extending in the xy-plane across the apical dendrite shaft (**Fig. 3a**, **Supplementary Fig. 6**).

**Fig 3.**
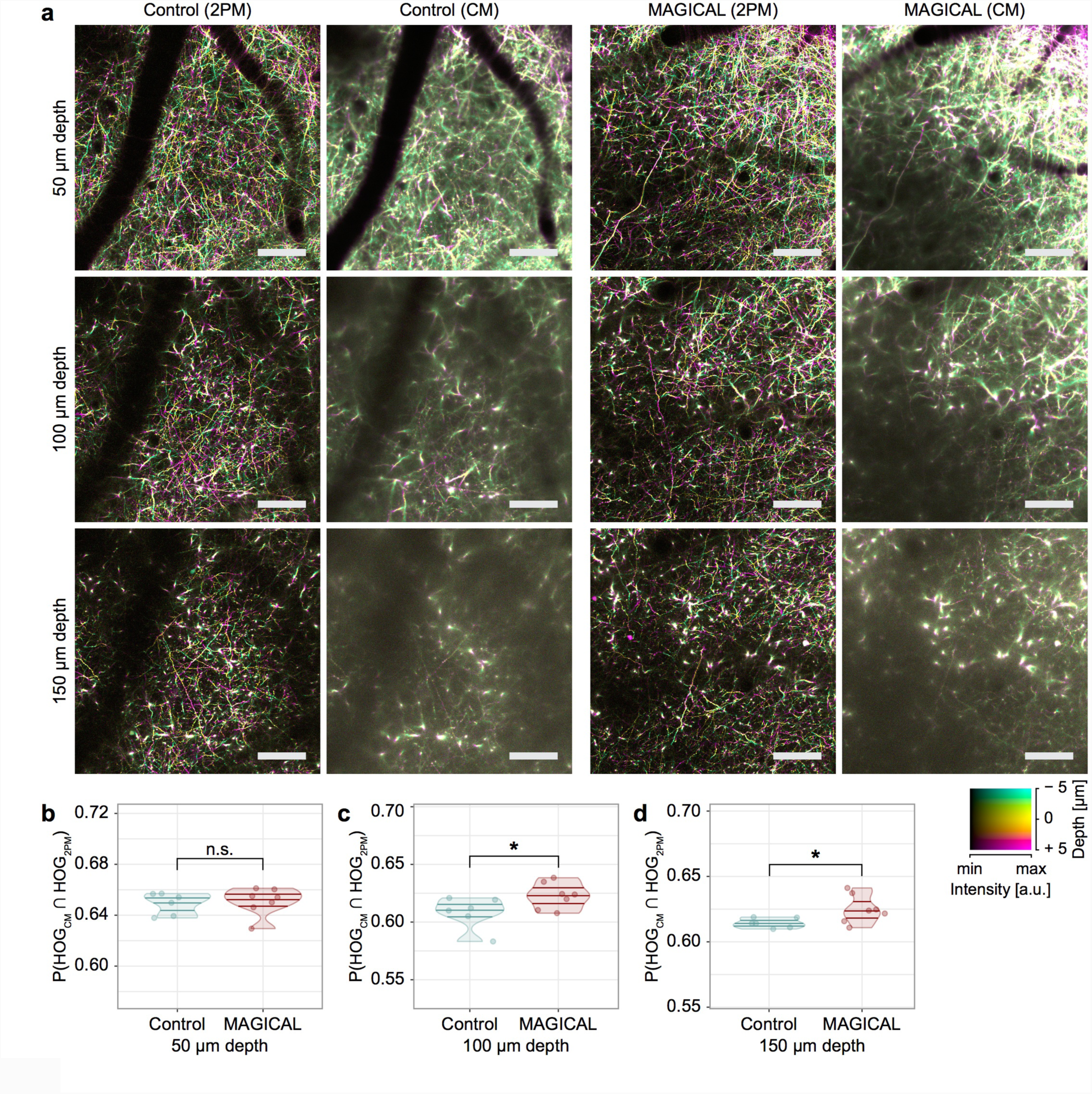
*In vivo* confocal imaging at CxLI. (**a**) DccMIP images of CxLI at 50 ± 5 μm, 100 ± 5 μm, and 150 ± 5 μm depth, by using two-photon microscopy (2PM) and confocal microscopy (CM). The display range of fluorescence intensity (min–max in 8 bits) is adjusted as fellow: 2PM images at every depth, (0–127); CM images at 50, 100, and 150 μm depth, (0–255), (0–127), and (0–63), respectively. In the color code, each center depth is shown as 0 μm relative depth. scale bar 100 μm. (**b–d**) Similarities of the DccMIP images between CM and 2PM at 50 ± 5 μm (**b**), 100 ± 5 μm (**c**), and 150 ± 5 μm (**d**) depth. P(HOG_CM_ ∩ HOG_2PM_), probability of intersection between HOG features extracted from the CM image and the 2PM image; *n* = 6 mice in control and *n* = 7 mice in MAGICAL; n.s., not significant; * *p* < 0.05; a.u., arbitrary unit.

We quantified similarities between CM images and 2PM reference images at same CxLI areas, by using the intersection probability between Histogram of Oriented Gradients (HOG) features extracted from each image. In the HOG comparison, CM images with MAGICAL retained more similarity to 2PM reference images at deeper regions (**Fig. 3b–d**, *p* = 0.0484 at 100 μm depth, *p* = 0.0420 at 150 μm depth, in permuted Brunner-Munzel test, *n* = 6 in control mice, *n* = 7 in MAGICAL mice). These results show that MAGICAL improves the observable depth at which *in vivo* confocal imaging can visualize detailed features.

To evaluate visualization abilities of confocal microscopy with MAGICAL, we captured 3D stack images with high-magnification at around 100 μm depth. In MAGICAL mice, the images showed synaptic structures, such as dendritic spines and axonal boutons, clearly, and fine fibers with less optical aberration along the z-axis (**Fig. 4**, **Supplementary Fig. 7**). Moreover, *in vivo* confocal images with MAGICAL did not show remarkable bleaching or phototoxic effects, as observed in time-lapse 3D stack imaging at 5-min intervals for 30 min (**Fig. 4**, **Supplementary Fig. 7**). These data suggest that MAGICAL would provide a practical approach to observe deeper regions with superior spatial resolution beyond the dilemma posed by confocal microscopy.

**Fig 4.**
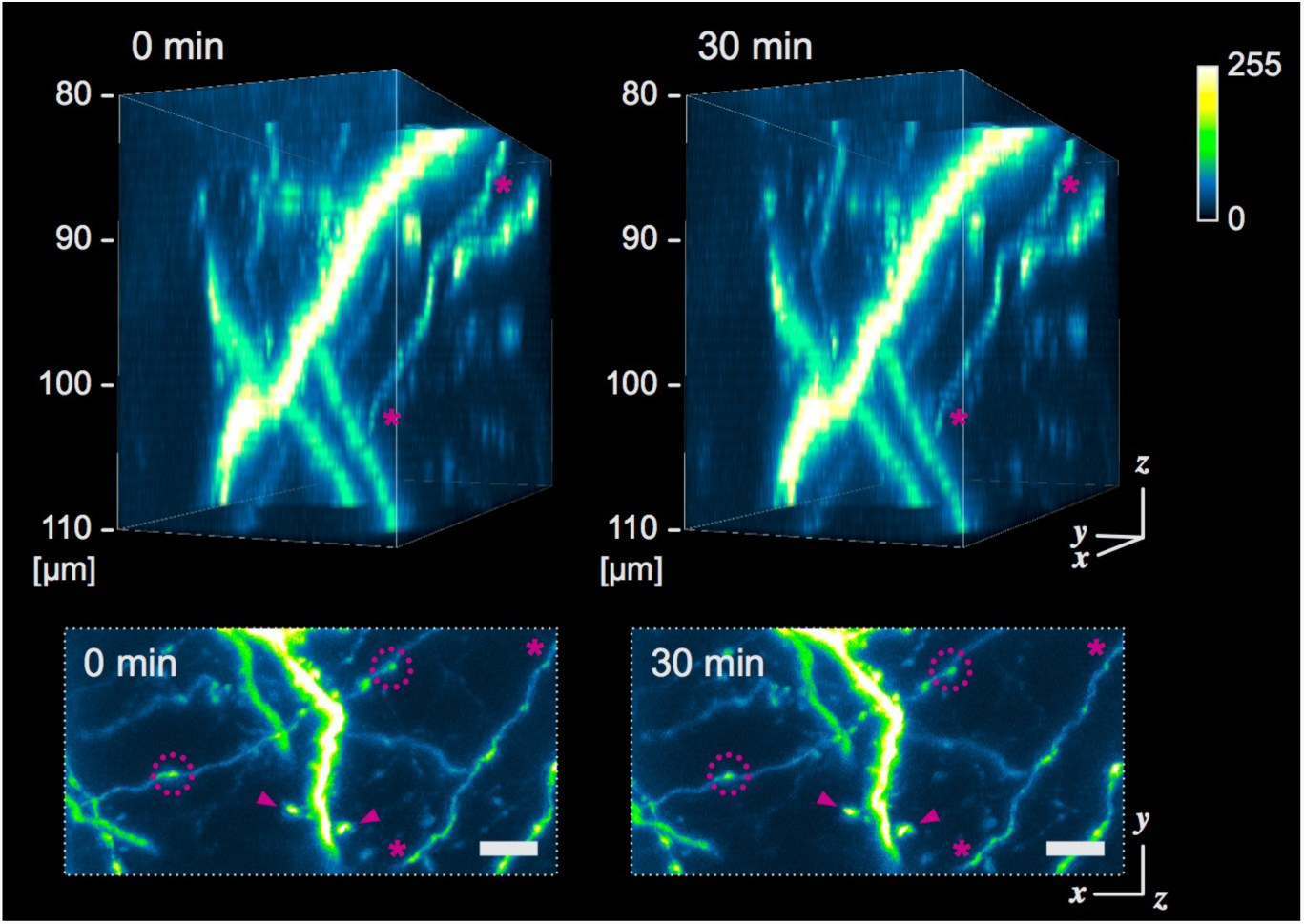
*In vivo* confocal high-resolution 4D imaging. Fine confocal 3D images with MAGICAL, captured with time lapse at 5-min intervals for 30 min. 3D images (upper) were reconstructed from 31 optical sections at 1 μm/step. A fine fiber (asterisk) was visualized with less optical aberration along the z-axis. In maximum intensity projection (MIP) images (bottom), synaptic structures, such as dendritic spines (arrow-head) and axonal boutons (dotted circle) were observable. Despite a totality of 217 scans, remarkable bleaching and phototoxic effects were not observed. scale bar 5 μm.

### Evaluation of clearing ability by FESTA

Fluorescence signal enhancement at several depths is one of the phenomena expected with optical clearing. However, it remains unclear whether MAGICAL improves transparency inside living brains. To address the question, we tried to evaluate *in vivo* transmittance at different wavelengths through the brain.

In conventional transmissive analyses, a certified light source is needed on the other side of a detector beyond a target, to calculate transmittance, i.e., the intensity ratio of transmissive light through a target to incident light from the source. In this perspective, fluorescence probes do not suit the light source for *in vivo* transmissive analysis, because original fluorescence intensity within focal point depends on the amount of the fluorescence probes and intensity of transmitted excitation light, both of which are unknown and uncontrollable inside living brains. However, a fluorescence emission spectrum is physically certified as emission probability at wavelengths, determined by energy levels and transition frequencies inherent in its molecule. Thus, to evaluate *in vivo* transmittance, we developed a new analytical method, Fluorescence Emission Spectrum Transmissive Analysis (FESTA), by using the emission spectrum as a scale independent of whole fluorescence intensity.

As fluorescence resources of *in vivo* FESTA, artificial yellow-green (YG) beads (Fluoresbrite YG Microspheres, Calibration Grade 1 μm; Polysciences) were injected into living mouse brains, and then were scattered in small clusters with cerebrospinal fluid flow. We calculated spectral intensity ratio of Cyan (458–511 nm) to Yellow (560–650 nm) light within *in vivo* images of the bead clusters (**Fig. 5a**). In contrast to fluorescence intensity, the Cyan/Yellow ratio was independent on both of amount of the fluorescence probes and two-photon excitation power (**Supplementary Fig. 8**). However, the Cyan/Yellow ratio was different between depths (**Supplementary Fig. 8**). Therefore, it is considered that the Cyan/Yellow ratio is proportional to a ratio, comprising the emission probability (constant), detector sensitivity (fixed as constant), and transmittance in the return path from inside the brain, for the Cyan and Yellow light.

**Fig 5.**
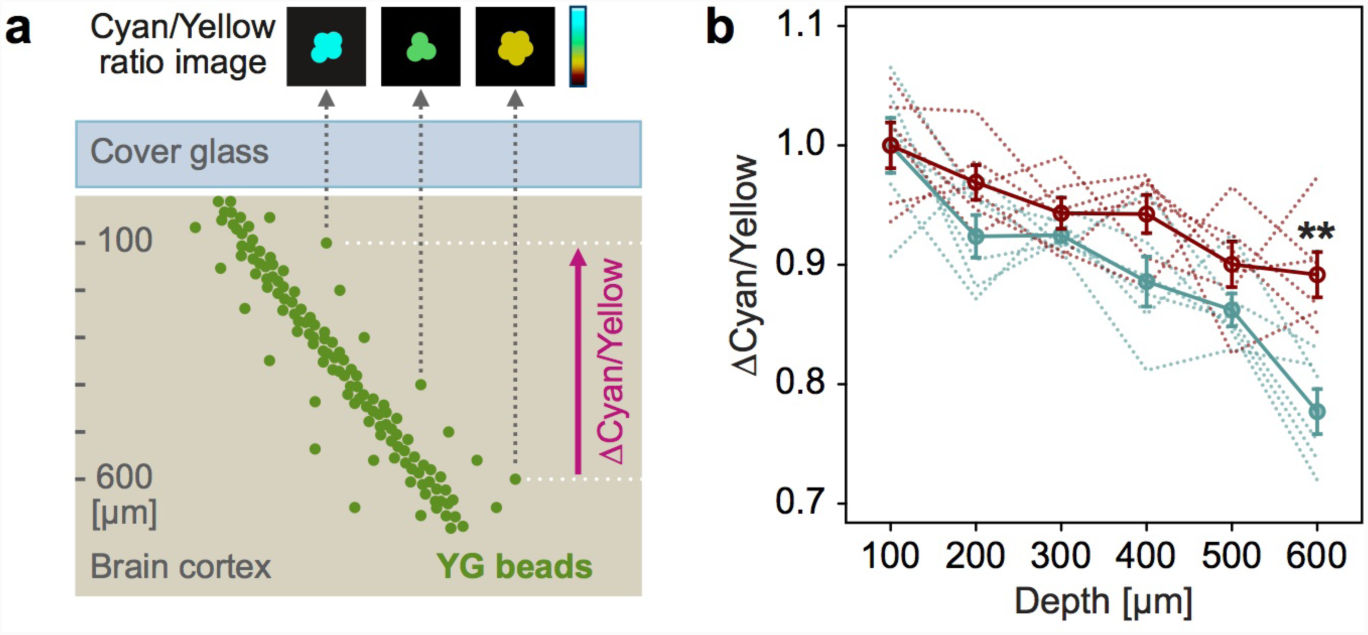
Optical evaluation for living brains. (**a**) Schematic illustration of the FESTA by using fluorescent YG bead clusters injected into living brains. (**b**) *In vivo* spectral transmission ratio (ΔCyan/Yellow) while the light returned from deeper regions to 100 μm depth. Red, MAGICAL; Blue, control; Solid line, average with standard error; Dotted line, each trial; *n* = 6 mice in each group; ** *p* < 0.01.

To focus on internal transmittance in living brains, and to remove the constant factors, we evaluated spectral transmission ratio (ΔCyan/Yellow) while the light returned from deeper regions to 100 μm depth. The ΔCyan/Yellow gradually decreased along the depth (**Fig. 5b**). It is, therefore, reasonable to consider that the cyan transmittance is more likely to be decreased than the yellow one because shorter wavelength light is more scattered depending on the optical path-length. However, in MAGICAL, a decrease of ΔCyan/Yellow was suppressed at deeper regions (*p* = 0.00159 in unpaired two-tailed Welch’s *t*-test, 95 % confidence interval (CI) = 0.0551–0.174, *t* = 4.29, *df* = 10.0, Cohen’s *d* = 2.48, *n* = 6 mice in each group, at 600 μm depth). These results suggest that MAGICAL improves *in vivo* transmittance especially of shorter wavelength light susceptible to optical scattering. Thus, we conclude that MAGICAL exerts an optical clearing ability to reduce scattering in living brains.

## Discussion

MAGICAL improved clear fluorescence signals, which would be enough to achieve faster imaging at deeper regions, *in vivo*. The imaging speed is one of the most important factors for *in vivo* observation. Fast imaging enables the visualization of not only instantaneous changes in neural activity with high temporal resolution but also finer structures of neural circuits by overcoming heartbeat-related trembles. In addition, fast imaging is required for enhancement of spatial resolution per time unit and expansion of observable areas within limited experimental time. Furthermore, fast imaging would be preferred to reduce phototoxic effects *in vivo*. Thus, MAGICAL would be a fundamental method for *in vivo* imaging to observe extensive neural circuits at high spatiotemporal resolution with less invasiveness.

Our results suggested that MAGICAL can transparentize living brains via scattering reduction. This leads us to the question of how does MAGICAL suppress the optical scattering in living brains? Generally, the major source of optical scattering in tissues is refractive index mismatches between structural components (n_d_; approx. 1.5) and interstitial fluids (water, n_d_; 1.33). A conventional optical clearing is achieved by increasing the refractive index of fluids with high index reagents, including glycerol (n_d_; 1.47 at 100 %), which can infiltrate into tissues and/or replace original fluids by itself, and as a result, reduce the mismatches. This clearing approach is useful for *in vivo* imaging to penetrate into tough tissues, such as skin^26^ and skull^27^, where glycerol immersion is tolerable even at high concentration keeping a high refractive index. However, glycerol also causes hemolysis *in vivo* as a function of dose, concentration, and route of administration^15,28^, indicating that the conventional technique using higher concentration glycerol is less applicable for almost *in vivo* tissues. Moreover, MAGICAL achieves optical clearing of living brains regardless of low concentration glycerol (5 %), which has almost the same refractive index as water. Therefore, to explain optical clearing by MAGICAL, other mechanisms different from conventional ones are needed.

MAGICAL was designed in reference to the medical treatment for cerebral edema and ischemic stroke^15-21^. Cerebral edema may occur for various reasons: cerebral trauma and inflammation allow the leak of serum proteins, drawing fluid into the brain due to the breakdown of the blood-brain barrier^29^. Furthermore, ischemia and hypoxia cause cell swelling due to a dysfunction of ion pumps caused by energy depletion^29^. The cerebral edema, in turn, reduces blood flow (ischemia) as a consequence of intracranial hypertension, thereby deteriorating hypoxia. Clinically, glycerol is commonly administrated to neurosurgery and stroke patients, in order to reduce edema via dehydration, thus to decrease intracranial pressure (ICP), and to increase cerebral blood flow^16,17,19-21^. From an optical point of view, light scattering is increased in living brains under cerebral ischemia and hypoxia, experimentally induced by blood removal and nitrogen gas inhalation, respectively^30,31^. The increased scattering is attributed to disturbances in cellular/subcellular structures due to the energy depletion. In the case of *in vivo* microscopy, living brains are exposed to loads, such as open-skull surgery, illumination for fluorescence excitation, and slight hypoxia from anesthesia. These loads cause brain trauma and inflammation, which in turn would trigger local edema, which, although not as severe as experimental ischemia and hypoxia, is enough to increase scattering for shorter wavelength light. Altogether, MAGICAL protects the living brains from edema and ischemia and thus would preserve the brain transparency.

What are the adverse effects of using MAGICAL for *in vivo* imaging? In toxicity tests, chronic oral glycerol has been shown as safe for animals, when administered in dosages equal to or lower than 9 g/kg-bw/day^32,33^. Thus, MAGICAL is considered as a non-invasive treatment, considering that 4.5-ml of 5 % (w/v) glycerol is daily administered as drinking water to a mouse weighing approximately 25-g. In the treatment of cerebral edema, glycerol administration does not show severe ICP rebound, electrolyte imbalance, and toxic side effects^16,17,20,21^. In addition, glycerol administration is expected to improve electroencephalogram and neurological status in ischemic stroke patients^17-19^. Moreover, glycerol administration is also commonly used to reduce intraocular pressure in glaucoma patients, without significant side effects suggestive of neurologic dysfunction^34^. These results suggest that MAGICAL would less disturb fundamental mechanisms executing brain functions. However, glycerol administration would improve energy metabolism in ischemic brains^19^, indicating that glycerol can be a cue to trigger some neural activity. The extent to which MAGICAL can modulate neural activity remains unclear, and thus probably should be evaluated for each neural circuit of interest.

MAGICAL is the first candidate for the optical clearing method applicable to living brains, easily improving *in vivo* microscopy without requiring any special skills or devices. Improved transparency for shorter-wavelength light with MAGICAL, means to expand the usability of various light sources, luminous probes, photosensitive caged compounds, and optogenetic tools for *in vivo* imaging and optical manipulation. In other words, MAGICAL is an important contribution toward for scientific advance, requiring a multi-laser/probe combination, such as the stimulated emission depletion microscopy^35^, photo-stimulation techniques^36^, and multi-color imaging^37^. Moreover, MAGICAL enables *in vivo* microscopy via the cranial window, to be combined with other neurological methods that insert electrodes or glass capillaries. These classical methods have been less compatible with *in vivo* microscopy due to an open-skull dogma that prohibits from contacting the brain for clear imaging. However, MAGICAL’s clearing ability has been demonstrated, despite after artificial beads injections in the FESTA. Thus, MAGICAL would accelerate functional analysis for brain functions, only by oral administration of glycerol.

## Methods

No statistical methods were used to predetermine sample size. In Generalized Linear Mixed Models (GLMMs), we used large sample sizes generally enough for modeling. For the Welch’s *t*-test, we confirmed that the *t*-test had adequate power, with a post-hoc test having effect size (Cohen’s *d*), *p*-value (< 0.01), and the sample size.

The experiments were not randomized. *In vivo* imaging must observe individual mice convergently, not parallelly, for a long time. To reduce influence dependent on specific timing, the observation order for each experimental group should be uniformly mixed. However, the sample size is insufficient for randomization to ensure to equalize the order. Thus, we allocated mice to experimental groups so as not to localize the order.

Investigators were not blinded to allocation during experiments and data analysis. To remove observer bias during experiments, observation area was selected by previously defined criteria, as described in “Methods, ‘Fluorescence intensity analysis’ section”. Data collection and analysis were automatically performed by software with fixed parameter setting or macro programs.

### Animals

All experiments were conducted in accordance with the Animal Research Committee of the Hokkaido University. The protocols were approved by the Committee on the Ethics of Animal Experiments in the Hokkaido University (No. 10-0119). Adult transgenic mice (9 to 12 weeks old, male) from Thy1-EYFP-H (H-line) were used for *in vivo* imaging of neurons. Adult C57BL/6NCrSlc mice (10 to 22 weeks old, male; Japan SLC, Shizuoka, Japan) were used for the FESTA with injected fluorescent beads. All mice were housed with food and drink *ad libitum* and were maintained in a 12 h light-dark cycle (lights on from 8:00 to 20:00), with controlled temperature (22 °C to 26 °C) and humidity (40 % to 60 %).

### Glycerol administration

For MAGICAL mice, 5 % (w/v) glycerol (075-00616; Wako Pure Chemical Industries, Osaka, Japan) was administered orally in the drinking water, *ad libitum*, from 2 weeks prior to open-skull surgery to the end of the experiment, except during operations and observations. Control mice received only water.

### Open-skull surgery

A cranial window was surgically created overlying the left cortex, according to a previous protocol^14,23^, with minor modifications. Briefly, 45 mg/kg-bw minocycline (Nichi-Iko Pharmaceutical, Toyama, Japan) were injected intraperitoneally (i.p.) about 4 h before surgery, to suppress bacterial infection and to protect neurons against microglial activation (neuroinflammation). Anesthesia was induced with an i.p. injection of 60 mg/kg-bw pentobarbital sodium (Somnopentyl; Kyoritsu Seiyaku Corporation, Tokyo, Japan), and was maintained with isoflurane inhalation (0.5 % to 1.5 %). To limit inflammation, 2 mg/kg-bw dexamethasone (Kyoritsu Seiyaku Corporation) was injected intramuscularly. Cranial bones were exposed, and then the left parietal bone was circularly carved about 4.2 mm in diameter with a dental drill and was removed gently. The exposed dura was washed with phosphate-buffered saline (PBS) to remove blood cells. A 4.2-mm diameter round coverslip (Micro Cover GLASS, #1S, about 0.17 mm thickness; Matsunami Glass Industry, Osaka, Japan), previously coated with a biocompatible polymer— Lipidure^®^ (CM5206E; NOF Corporation, Tokyo, Japan), to prevent foreign-body reactions, especially to blood clotting—was placed directly over the dura without extra pressure, and was sealed on the bone edges with cyanoacrylate cement and dental adhesive resincement (Super-Bond C&B; Sun Medical Company, Shiga, Japan). To make the foundations for chamber fixation, exposed cranial bones around the cranial window were partially coated with the adhesive resin cement. A head chamber, which has a center hole to expose the cranial window, was secured to the cranial bones with dental acrylic resin cement (Unifast III; GC Corporation, Tokyo, Japan). 5 mg/kg-bw carprofen (Rimadyl; Zoetis, Parsippany, NJ, USA) was injected i.p. to reduce inflammation and pain. After surgery, the mouse was singly housed for recovery at least 3 hours until observation.

### Beads injection

The beads injection into the cortex was performed in an open-skull surgery, with minor modifications before sealing with the coverslip, by using an oil filled glass micropipette attached to a mechanically driven Hamilton microsyringe. The left parietal bone was elliptically resected about 3.9-mm in the anterior-posterior (A-P) axis and about 2.7-mm in the medial-lateral axis. A needle of the glass micropipette was inserted into the cortex at an angle of 54 degrees with respect to the brain surface, and proceeded 0.9 mm towards the anterior direction along the A-P axis, to reach about 0.72 mm depth. 1-μm YG beads (Fluoresbrite^®^ YG Microspheres, Calibration Grade 1 μm, #18860; Polysciences, Incorporated, Warrington, PA, USA) were diluted at 4.6 × 10^8^ beads/mL with PBS, and were gradually injected into the needle trace in repeated steps, in which the needle was withdrawn 0.3 mm and then was kept in place for 5 min. During and after the injection, the exposed dura was washed with PBS to remove blood cells and to prevent drying. After the surgery, the mouse was singly housed for recovery at least one day until observation.

### Setup of *in vivo* imaging

All *in vivo* imaging sessions were conducted in mice anesthetized with isoflurane inhalation^23,24^. The head chamber was mounted and glued to a customized adaptor stage, which suspended the mouse body with a harness, to reduce adverse effects due to movements. After setting under an upright microscope system, the adaptor stage was adjusted at a tilt angle, referring to 1-μm diameter fluorescent beads scattered on the coverslip as a guide, so that the cranial window was positioned horizontally, hence providing a suitable optical alignment. All *in vivo* imaging sessions were performed under a microscope system (A1R MP+ Multiphoton Confocal Microscope; Nikon, Tokyo, Japan) controlled by NIS-Elements software (version 4.13.00; Nikon) with 12-bit dynamic range. Before *in vivo* imaging, z-position at the brain surface was set as zero for absolute depth.

### Two-photon microscopy

*In vivo* two-photon imaging was performed with a water immersion objective (25× Ob., 1.10 NA) in PBS. The EYFP and the YG beads were excited by 960 nm and 900 nm, respectively, emitted from Ti:Sapphire laser (Mai Tai DeepSee; Newport Spectra-Physics, Irvine, CA, USA). By using a Galvano scanner in one-directional mode, the xy-images in the 3D stacks were acquired as 512 × 512 pixels at 1 frame per sec (fps), 2.2 μsec per pixel, which is maximum speed in this condition. The 3D display was performed using the NIS-Elements software.

To visualize the neural circuits, all the EYFP signals under 650-nm wavelength were collected in a GaAsP type Non-Descanned Detector (GaAsP-NDD). The CxLV and HpCA1 images were captured as 3D stacks with 3 μm z-step size. The depth of the CxLV was determined between 495 μm and 585 μm, where somas of CxLV pyramidal neurons were observed in every mouse. The depth of the HpCA1 was slightly different for each mouse, thus the middle of stratum pyramidale, approximately at 975 μm absolute depth, was set as the center depth of each HpCA1 stack. To evaluate the brightness of the images, laser power (LP) under objective lens and detector sensitivity (HV: high voltage) were fixed. The CxLV was captured with LP 66 mW (under 25× Ob.) and HV 30. The HpCA1 was captured with LP 200 mW (under 25× Ob.) and HV 30.

For the FESTA, fluorescence signals were split to multi-colors by dichroic mirrors (DM) in front of the GaAsP-NDDs. The signals of YG beads were split by DM458, DM511, and DM560, and then separately detected at 458–511 nm (Cyan) and at 560–650 nm (Yellow). To avoid optical influence by injection trace, the FESTA applied to the beads clusters observable without crossing the trace. The beads images were captured as 3D stacks with 0.25 μm z-step size at depth of interest ± 1.0 μm (9 sections). The detector sensitivities were fixed: HV 1 on the Cyan channel and HV 3 on the Yellow channel. The laser powers were adjusted for each bead cluster, because bead clusters show different brightness depending on cluster size and injection depth.

### Confocal microscopy

*In vivo* confocal imaging was performed with a water immersion objective (25× Ob., 1.10 NA or 60× Ob., 1.20 NA) in PBS. The EYFP was excited by 488 nm, and its signal was collected at 500–550 nm in a standard detector. By using a Galvano scanner in one-directional mode, the xy-images comprising 3D stacks were acquired as 512 × 512 pixels at 1 fps or 512 × 256 pixels at 2 fps, 2.2 μsec per pixel, which is maximum speed in this condition.

To evaluate observable depth in confocal imaging, the CxLI images were captured as 3D stacks with 1 μm z-step size at 50 ± 5, 100 ± 5, and 150 ± 5 μm depth (11 sections) via a 25× Ob. Laser power and detector sensitivity were fixed at 10 % and HV 90, respectively. As references for confocal imaging, the same 3D stacks were also captured by using a two-photon microscope, of which detector sensitivity was fixed at HV 30, but 960-nm laser powers were adjusted to 12.5, 16.1, and 19.2 mW (under 25× Ob.) at 50, 100, and 150 μm depth, respectively.

High-resolution 4D imaging with a 60× Ob. was performed as 3D stacks with 1 μm z-step size at approximate 100 ± 15 μm depth (31 sections), as a time-series at 5 minutes intervals for 30 minutes (7 time-points). Laser power and detector sensitivity were set between 10 and 20 % and between HV 90 and HV 100, respectively. The 3D display was performed by NIS-Elements software.

### Fluorescence intensity analysis

For two-photon imaging at CxLV and HpCA1, 30 mice (Treatment: control, *n* = 15; MAGICAL, *n* = 15) were observed. For confocal imaging at CxLI, 17 mice (Treatment: control, *n* = 8; MAGICAL, *n* = 9) were observed. In the H-line mice, signal distribution is uneven, especially at the somatic layer, because the EYFP expression level varies among positive neurons, which tend to be differently clustered in different individuals. To avoid selection bias due to individual differences, the imaging area was selected according to the xy position where the hippocampus was observable at the most shallow depth.

Fluorescence intensity analysis was performed on the 3D image stacks, by using Fiji/ImageJ software (version 2.0.0-rc-49/1.51d) ^38^. For sorting of image stacks within each capture condition, mice were ranked in descending order of brightness (average fluorescence intensity per pixel) extracted from image stacks around the center region: at 540 ± 12 μm for the CxLV, approximately at 975 ± 12 μm for the HpCA1, and at 100 ± 5 μm for the CxLI. The top or bottom brightness at each capture condition came from over-or under-exposed image stacks, respectively. Thus, mice having the top or bottom brightness within each treatment group were equally removed from the following analysis, as outliers of the capture condition. Finally, 26 mice (control, *n* = 13; MAGICAL, *n* = 13) and 13 mice (control, *n* = 6; MAGICAL, *n* = 7) were evaluated in fluorescence intensity analysis for two-photon imaging and confocal imaging, respectively. A representative image for each treatment group at each capture condition was obtained from the mouse scored as intermediate brightness.

In the two-photon imaging, we evaluated the fluorescence intensity distributions (FIDs) under HV 30, at three different depths within the observation area. For the CxLV, the three depths were set at 510 ± 3, 540 ± 3 (center), and 570 ± 3 μm depth. For the HpCA1, three depths were relatively adjusted at the middle of stratum pyramidale as the center (0 μm), approximately at 975 ± 3 μm absolute depth, and at 24 μm above and below the center. These three depths were sufficiently far away from each other to ensure separate images in the mice.

The FIDs were summarized to 8-bit intensity bins histograms by the Fiji and were visualized through following steps by R (version 3.3.2) ^39^ with RStudio software (version 1.0.44). The histograms were arranged as a matrix, where the image stacks were sorted in rows according to their brightness in decreasing order, and the intensity bins were aligned in columns. Pixel counts in the histogram were converted with log_10_(z+1), where 1 is a constant to avoid log_10_(0) contamination in the converted matrix. The matrix of FIDs was visualized as a heat map. Pixel counts (excluding zero) were ranked in percent quantile. In the heat maps, the 50 % quantile was shown as cyan dotted contours.

### Image similarity analysis

Based on image features, similarities between CM images and 2PM reference images at same CxLI areas were evaluated in 13 mice (Treatment: control, *n* = 6; MAGICAL, *n* = 7), which passed fluorescence intensity analysis to remove outliers. The 3D stack images were converted to DccMIP 2D images with a color code shown in **Fig. 3a**, by using Fiji/ImageJ software. The display range of fluorescence intensity (min–max in 8 bits) were adjusted as fellow: 2PM images at every depth, (0–127); CM images at 50, 100, and 150 μm depth, (0–255), (0–127), and (0–63), respectively.

Features extraction and comparison were performed by using OpenCV (version 3.4.2) via a python (version 3.6.6) code. In gray-scale space, a Histogram of Oriented Gradients (HOG) ^40^ feature was extracted from each image (512 × 512 pixels) with parameters as fellow: window size, (512, 512); block size, (128, 128); block stride, (4, 4); cell size, (16, 16); nbins, 180; other parameters were used as preset in OpenCV. The raw HOG feature comprising 108,391,680 bins was normalized to 1 to convert a probability histogram. In the comparison between paired two images, a similarity index was defined as the intersection probability of the HOG features.

### Fluorescence Emission Spectrum Transmissive Analysis

For spectral ratio calculation, data processing was performed by using the Fiji/ImageJ software (version 2.0.0-rc-49/1.51d) ^38^. All xy images in raw data (multi-color 3D stack) were smoothed with the median filter (1 pixel). Pixels having faint or saturated signal values (< 32 or 4063 < in 12 bit, respectively) were excluded from the following calculation. Background area out of beads was detected by an auto-threshold function with the Otsu method on the Yellow channel and was removed from all channels. Spread bead cluster images were selected by ROI (region of interest) to exclude the injection trace from the calculation. The pixels in YG bead clusters were plotted on a scatter graph, corresponding to the Yellow intensity along the x-axis and the Cyan intensity along the y-axis. At depth of interest, representative Cyan/Yellow ratio was determined from the slope of the best-fitted regression line (highest *R*^2^) on the scatter plot, derived from image stacks at its depth ± 1.0 μm. To calculate the ΔCyan/Yellow, the Cyan/Yellow ratio was divided by the average of Cyan/Yellow within the experimental group at 100 μm depth.

### Statistics

All statistical analysis was performed by using R (version 3.3.2) ^39^ with RStudio software (version 1.0.44). Conventional two-tailed Welch’s *t*-test and Cohen’s *d* were calculated with the default and the MBESS package (version 4.3.0), respectively. Permuted Brunner-Munzel^41-43^ test was executed by the lawstat package (version 3.1) with permutation code.

To assess MAGICAL effects on FIDs, we constructed GLMMs with a Gamma error distribution and a log link function. In the models, fluorescence intensities of pixels (FIPs) were used as a response variable. The FID of each image stack at the depth ± 3 μm, therefore, was re-expanded to an intensity list comprised of 786,432 pixels (512 × 512 pixels × 3 sections). To reduce modeling costs, 12,288 pixels (1.56 %) were randomly sampled without overlap from each list. Saturated pixels (255 in 8 bit) were excluded from modeling due to inaccuracies in fluorescence intensity. Zero intensity pixels were also excluded to avoid a calculation failure under the log-link function. “Treatment” (2 levels, MAGICAL or control) and “Depth” (3 levels) were used as categorical predictor variables with fixed effects. The models included an interaction between “Treatment” and “Depth.” To overcome pseudo-replications caused by sampling multiple pixels within each mouse, “Mouse” (26 mice in total) was assigned as a random effect for intercepts and “Depth” slopes. Thus, the models were described in R as follows: FIP ∼ Depth * Treatment + (Depth | Mouse) Consequently, the models comprised 929,331 (CxLV) or 957,264 (HpCA1) pixel observations, in 78 image stacks extracted at 3 depths from 26 mice under 2 treatments.

The model fitting was performed via the maximum likelihood according to Laplace approximation, by using the glmer function in the lme4 package (version 1.1-12). For the estimated coefficient of fixed effect, Wald-type 95 % confidence interval (CI) was calculated by using the confint function. The model prediction for each image stack was obtained through the predict.merMod function in the lme4 package. The fixed effects with asymptotic 95% CI were estimated by using the Effect function in the effects package (version 3.1-2). Pairwise comparisons for the predictor variables were examined by the lsmeans function in the lsmeans package (version 2.25-5).

As the coefficient of determination for the GLMM, two *R*^2^ statistics are proposed^44^: marginal and conditional *R*^2^_*GLMM*_. The marginal *R*^2^_*GLMM*_ (*R*^2^_*GLMM(m)*_) describes the proportion of variance explained only by fixed effects. The conditional *R*^2^_*GLMM*_ (*R*^2^_*GLMM(c)*_) describes the proportion of variance explained by both fixed and random effects.

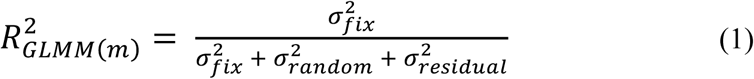

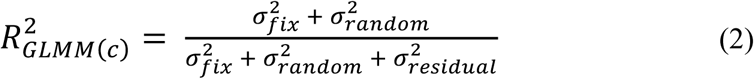

An original method for the *R*^2^_*GLMM*_ was limited only to the GLMM including Poisson or binomial error distributions and not including random slope terms^44^. However, two extensions had been individually provided for the *R*^2^_*GLMM*_, to calculate random effects variances in random slope models^45^ or to evaluate further error distribution in the residual variance^46^. Thus, we calculated the *R*^2^_*GLMM*_ of our models by an R code combining the two extensions as follows: in the extension for random slope models^45^, provided code generates a square matrix at once but uses only diagonal elements of the matrix. When the code ran for our models, a large number of observations generated a huge square matrix, which caused the computation to fail, due to insufficient memory. Therefore, we calculated only the portion corresponding to the diagonal of the square matrix. On the other hand, the extension for residual variances was applicable to our models without modifications. We selected the trigamma function from three candidates in the paper^46^, to evaluate the models including a Gamma error distribution and a log link function.

## Data availability

Data and code are available from the authors upon request.

## Endnotes

### Acknowledgments

We thank Nikon Imaging Center (NIC) at Hokkaido University for technical support with a 60× objective lens. This research was supported by the “Brain/MIND” from the Japan Agency for Medical Research and Development (AMED); by the MEXT/JSPS KAKENHI Grant Number JP15H05953 “Resonance Bio” from the Japan Society for the Promotion of Sciences (JSPS) in the Ministry of Education, Culture, Sports, Science and Technology (MEXT); by the “Network Joint Research Center for Materials and Devices” in the MEXT and by the “Dynamic Alliance for Open Innovation Bridging Human, Environment and Materials” in the MEXT.

## Author Contributions

K.I., R.K., and T.N. designed the research. K.I. and T.O. performed experiments and analyzed the data. K.I. and T.N. wrote the paper.

## Additional Information

The authors declare no conflicts of interest related to this research. Correspondence and requests for materials should be addressed to T.N.

