## Supplementary Information for "Optical clearing of living brains with MAGICAL to extend *in vivo* imaging"

**Title:**

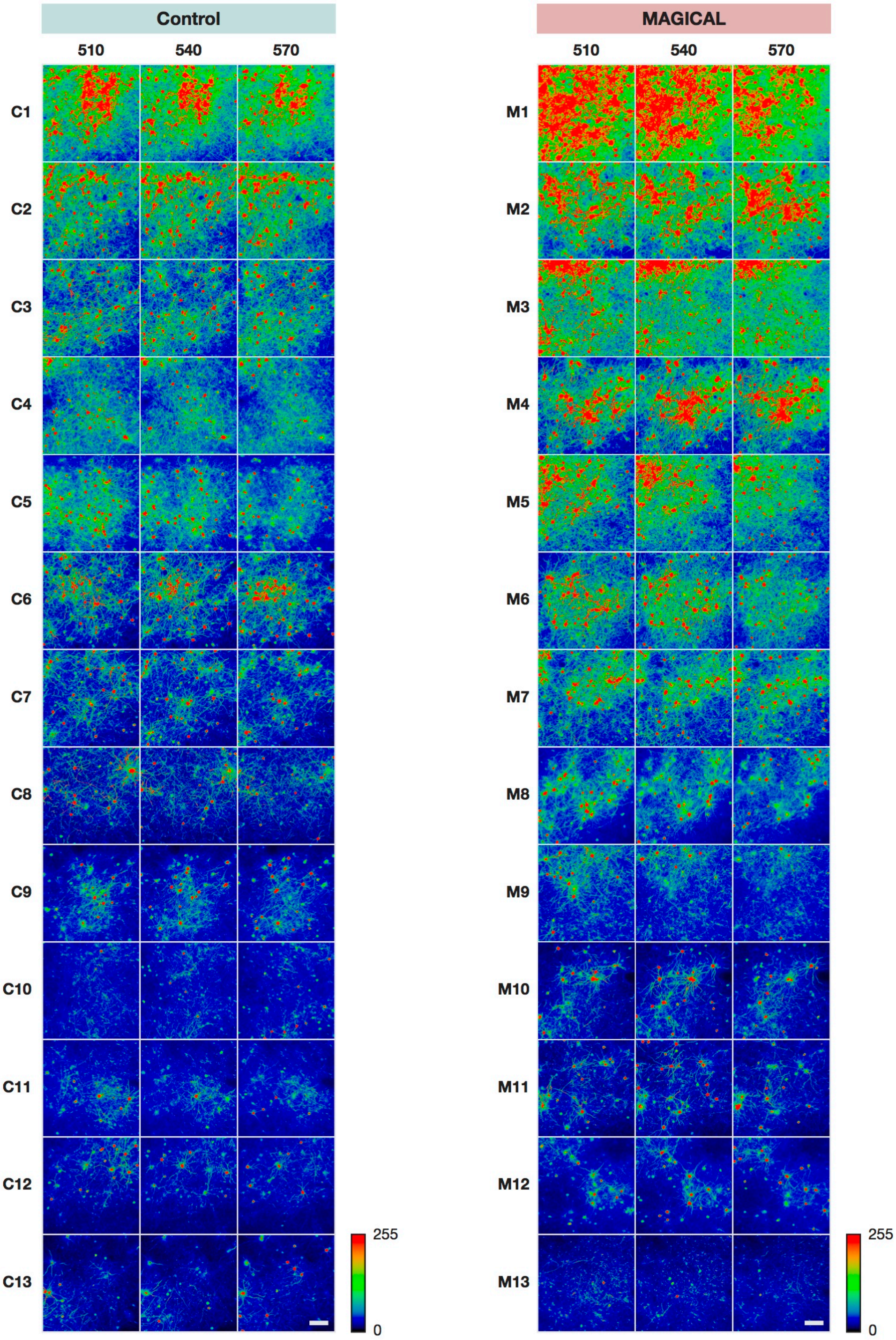


**Supplementary Figure 1. *In vivo* two-photon images at CxLV to evaluate fluorescence intensity.** All images were captured under the 25× Ob. with LP 66 mW and HV 30, as 3D stacks at indicated depth ± 3 µm with 3-µm z-step size (3 sections). These 3D stacks were analyzed as FIDs (**Supplementary Fig. 2**) and FIPs for the GLMM (**Fig. 1b–c**). All 3D stacks are displayed as MIP images, ranked by brightness scores (average fluorescence intensity per pixel) at 540 ± 12 µm depth (9 sections) captured with LP 66 mW and HV 30. scale bar 100 µm.

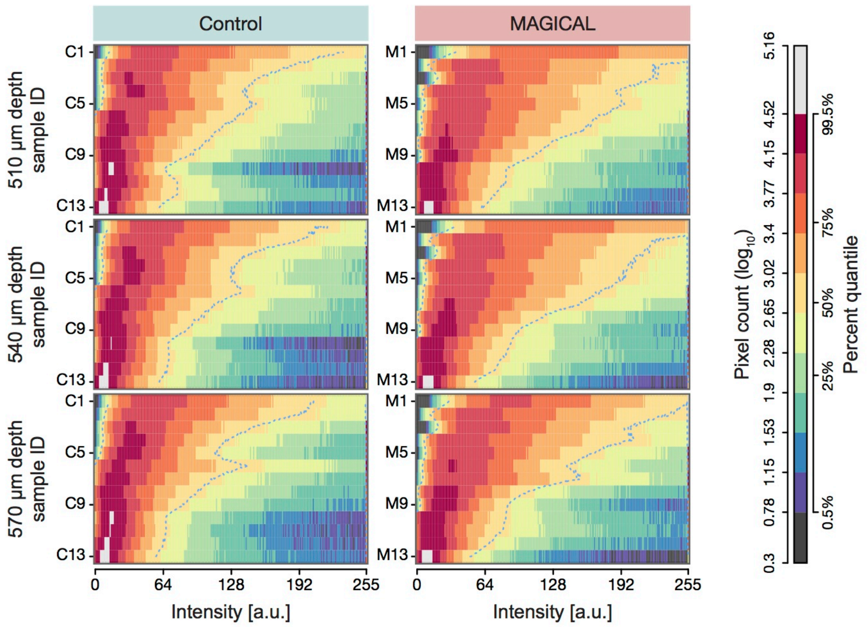


**Supplementary Figure 2. Heat map representation for FID of CxLV image stacks in Supplementary Fig. 1.** The heat map rows represent image stacks ranked by brightness scores at 540 ± 12 µm depth within each experimental group. The heat map columns represent intensity bins (8 bits). Pixel counts (z) in the FID histogram are converted with $\log_{10}(z+1)$ and assigned a color code. Percent quantile ranks non-zero pixel counts. Cyan dotted contours are median. a.u., arbitrary unit.

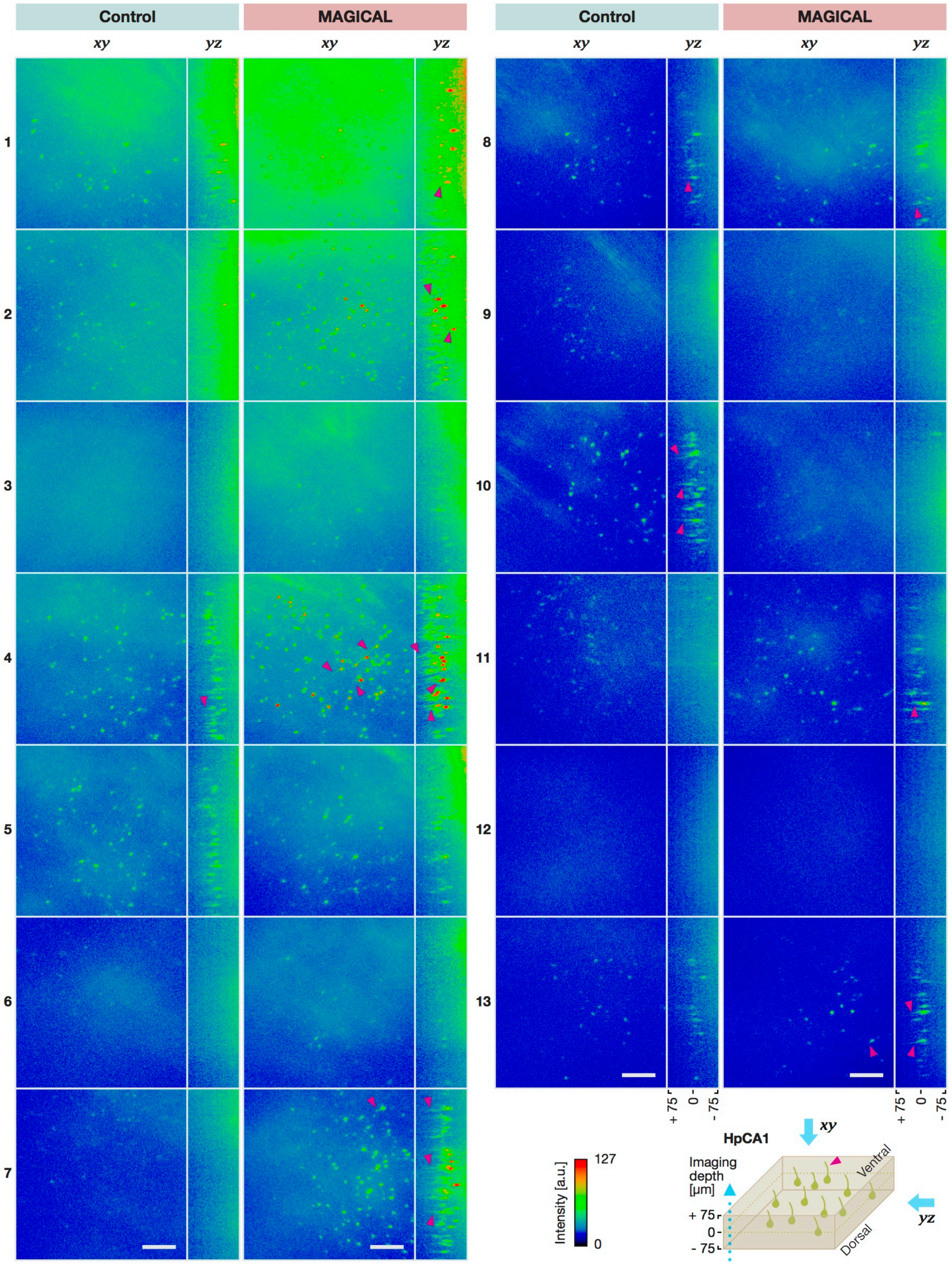


**Supplementary Figure 3. *In vivo* two-photon 3D rendering images at HpCA1.** All images were captured under the 25× Ob. with LP 200 mW and HV 30, as 3D stacks at the middle of stratum pyramidale (0 μm relative depth) ± 75 μm with 3-μm z-step size (51 sections). The 3D rendering was performed by using the Fiji's 3D project function with the brightest point mode and 50 % interior depth-cueing. Xy projection images were created from a ventral viewpoint (opposite side of observation), to avoid halation on the alveus (fibrous cloud structures) at the dorsal side. Yz projection images were created with rendering interpolation. These images are ranked by the brightness scores at 0 (center) ± 12 μm depth (9 sections). MAGICAL visualized many tiny apical dendrites (arrow-head) of pyramidal cells, although it is dependent on the signal to the background ratio. scale bar 100 μm. a.u., arbitrary unit.

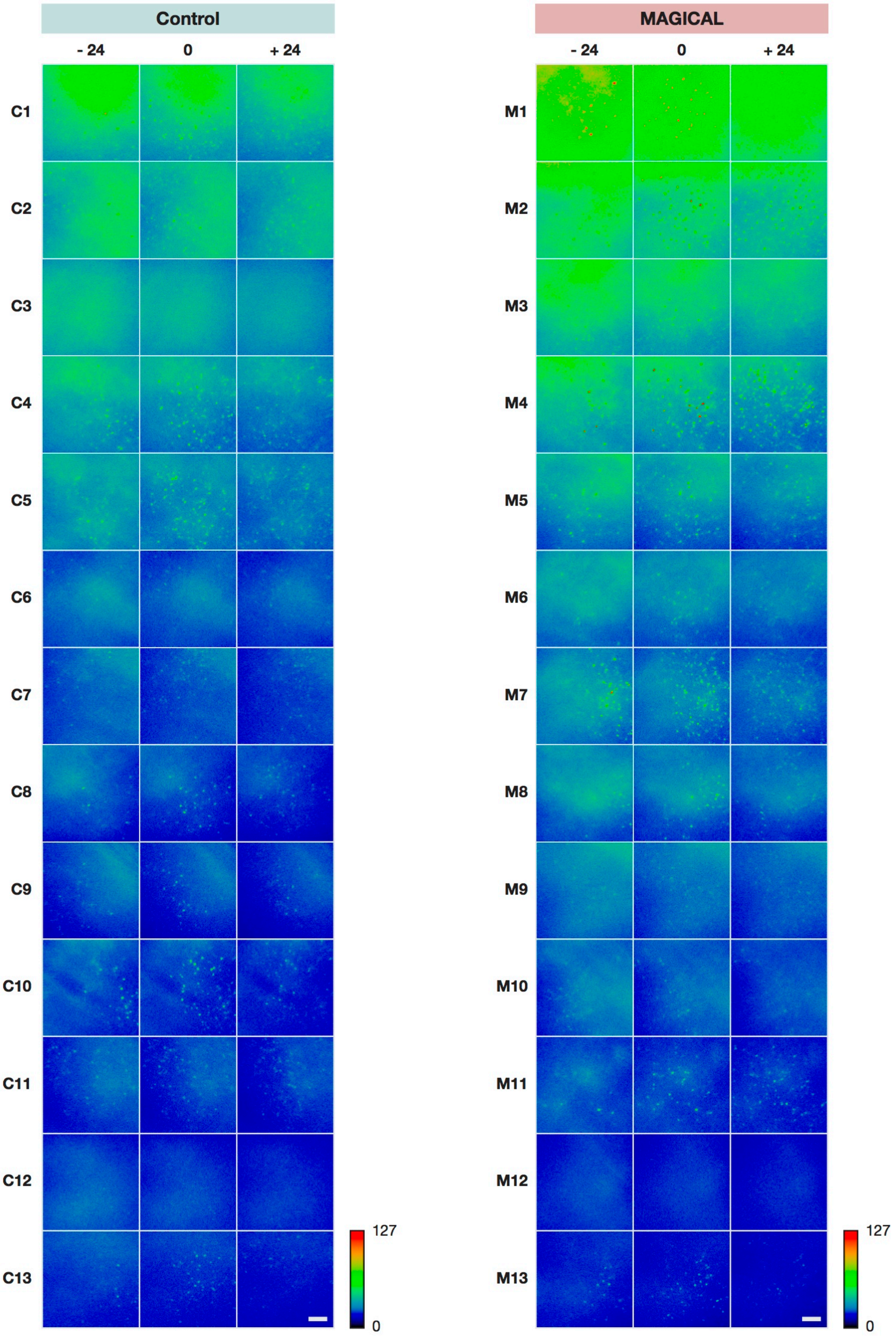


**Supplementary Figure 4. *In vivo* two-photon images at HpCA1 to evaluate fluorescence intensity.** All images were captured under the 25× Ob. with LP 200 mW and HV 30, as 3D stacks at indicated relative depth ± 3 μm with 3-μm z-step size (3 sections). These 3D stacks were analyzed as FIDs (**Supplementary Fig. 5**) and FIPs for the GLMM (**Fig. 2b–c**). All 3D stacks are displayed as MIP images, ranked by brightness scores at 0 (center) ± 12 μm depth (9 sections) captured with LP 200 mW and HV 30. scale bar 100 μm.

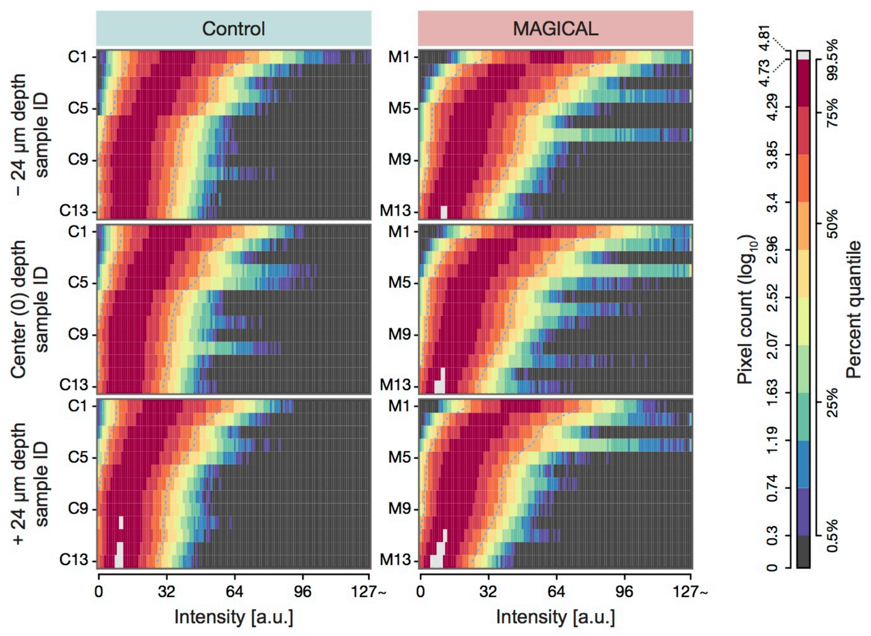


**Supplementary Figure 5. Heat map representation for FID of HpCA1 image stacks in Supplementary Fig. 4.** The heat map rows represent image stacks ranked by brightness at the center ± 12 µm depth within each experimental group. The heat map columns represent intensity bins (8 bits). Pixel counts (z) in the FID histogram are converted with $\log_{10}(z+1)$ and assigned a color code. Percent quantile ranks non-zero pixel counts. Cyan dotted contours are median. a.u., arbitrary unit.

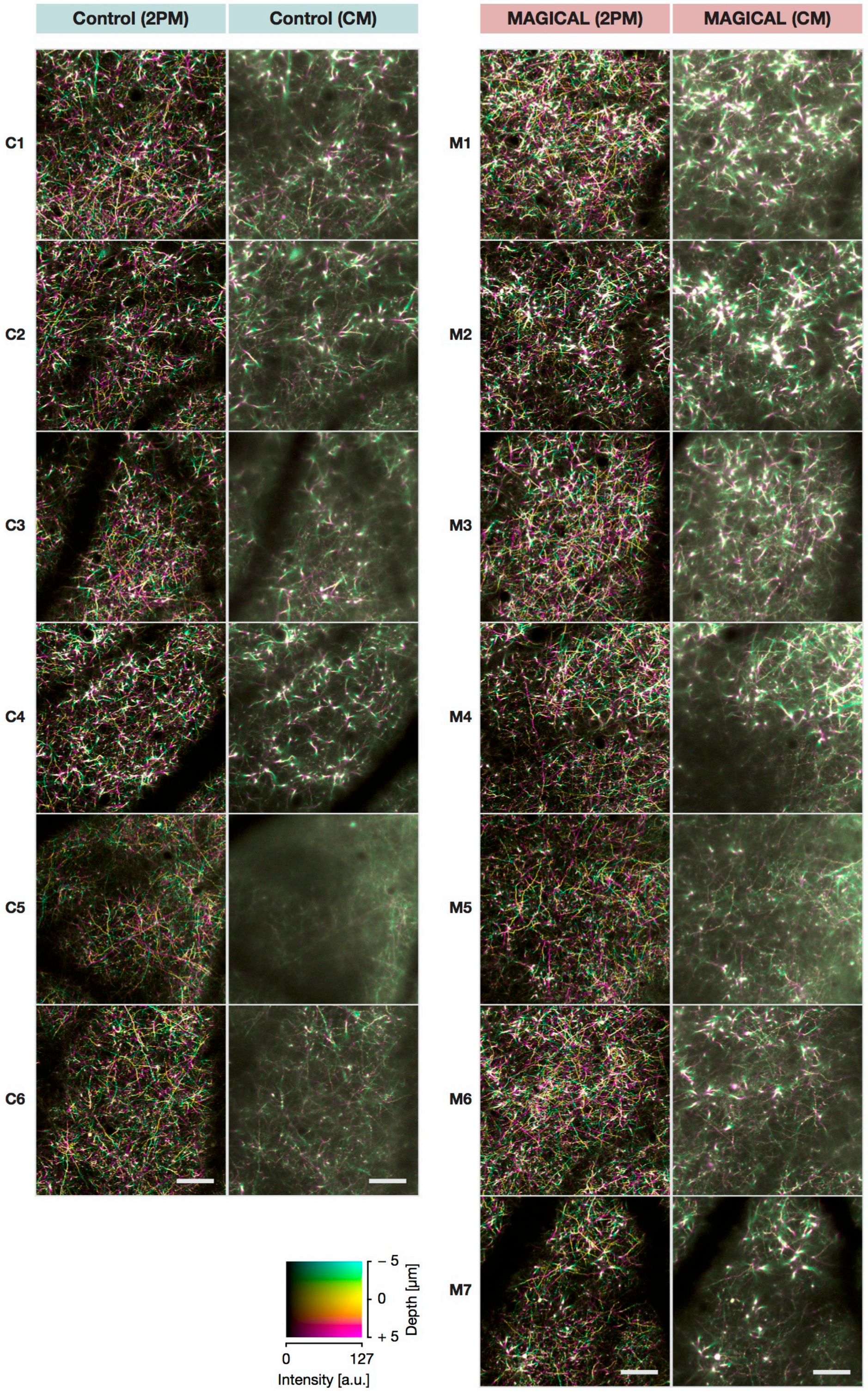


**Supplementary Figure 6. *In vivo* confocal images at CxLI.** All images were captured as 3D stacks at 100 μm (center, 0 μm relative depth) ± 5 μm with 1-μm z-step size (11 sections) under the 25× Ob., with LP 10 % and HV 90 in confocal microscopy and LP 16.1 mW and HV 30 in two-photon microscopy. All 3D stacks are displayed as DccMIP images, ranked by brightness scores in the confocal stacks. Intermediate images (C3 and M4) are shown as the representative images in **Fig. 3a**. scale bar 100 μm. a.u., arbitrary unit.

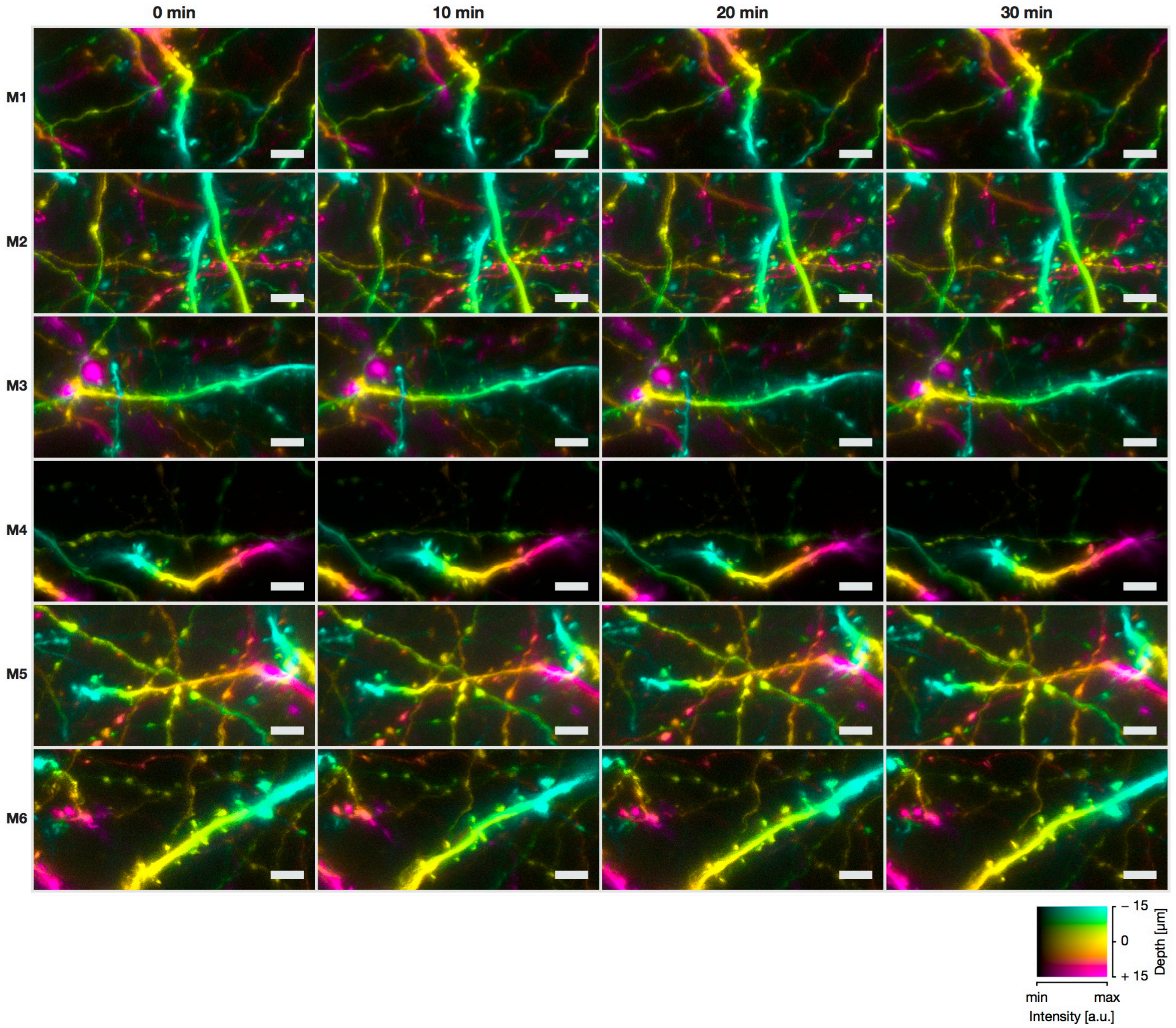


**Supplementary Figure 7. *In vivo* confocal high-resolution 4D images at CxLI.** In 6 mice with MAGICAL, the 4D imaging was performed as 3D stacks with 1 µm z-step size at approximate 100 ± 15 µm depth (31 sections, displayed as DccMIP), as a time-series at 5 minutes intervals for 30 minutes (7 time-points, but images at 5, 15, and 25 min were not shown). Laser power and detector sensitivity were set between 10 and 20 % and between HV 90 and HV 100, respectively. In the color code, each center depth is shown as 0 µm relative depth. scale bar 5 µm. a.u., arbitrary unit.

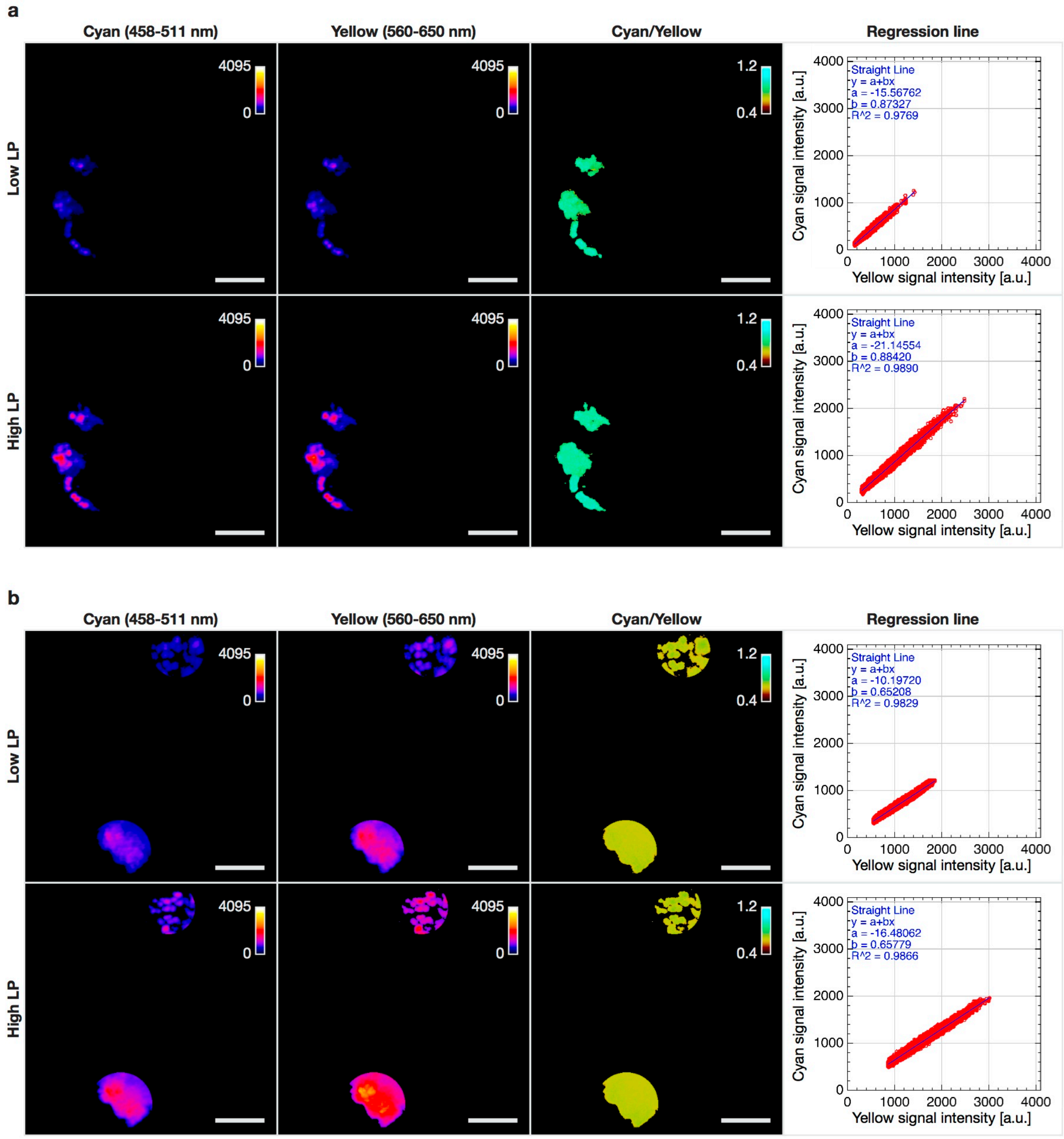


**Supplementary Figure 8. Cyan/Yellow ratio calculation for FESTA.** (**a–b**) MIP images of beads clusters extracted via ROI, and respective regression lines at 100 µm (**a**) and 600 µm (**b**) depth. Although fluorescence intensity in each channel was affected by LP for two-photon excitation and amount of fluorescence probe dependent on accumulation style of round-shaped beads, the Cyan/Yellow ratio was not affected. In contrast, the Cyan/Yellow ratio decreased with depth. scale bar 10 µm. a.u., arbitrary unit.

### Supplementary Table 1. The linear predictors in the GLMM.

**a**

Fixed Effects :

| Coefficient | Estimate | 95 % CI lower | 95 % CI upper | Std. Error | t value | Pr ( > \|z\| ) |
|---|---|---|---|---|---|---|
| (Intercept) | 3.595 | 3.522 | 3.668 | 0.037 | 96.92 | < 0.001 |
| Depth_540 | -0.024 | -0.049 | 0.000 | 0.013 | -1.92 | 0.0548 |
| Depth_570 | -0.044 | -0.080 | -0.009 | 0.018 | -2.43 | 0.0153 |
| Treatment_MAGICAL | 0.259 | 0.164 | 0.355 | 0.049 | 5.33 | < 0.001 |
| Depth_540 : Treatment_MAGICAL | 0.010 | -0.024 | 0.044 | 0.017 | 0.57 | 0.5710 |
| Depth_570 : Treatment_MAGICAL | -0.032 | -0.080 | 0.016 | 0.024 | -1.30 | 0.1928 |

Random Effects :

| Group | Name | Variance | Std.Dev. |
|---|---|---|---|
| Mouse | (Intercept) | 0.205 | 0.452 |
| | Depth_540 | 0.002 | 0.050 |
| | Depth_570 | 0.006 | 0.077 |
| Residual | | 0.649 | 0.806 |

Number of observations: 929331, groups: Mouse, 26

**b**

Fixed Effects :

| Coefficient | Estimate | 95 % CI lower | 95 % CI upper | Std. Error | t value | Pr ( > \|z\| ) |
|---|---|---|---|---|---|---|
| (Intercept) | 3.044 | 2.973 | 3.115 | 0.036 | 84.31 | < 0.001 |
| Depth_000 | -0.114 | -0.122 | -0.105 | 0.004 | -27.01 | < 0.001 |
| Depth_+24 | -0.230 | -0.244 | -0.217 | 0.007 | -33.43 | < 0.001 |
| Treatment_MAGICAL | 0.160 | 0.086 | 0.233 | 0.037 | 4.26 | < 0.001 |
| Depth_000 : Treatment_MAGICAL | -0.008 | -0.020 | 0.004 | 0.006 | -1.35 | 0.177 |
| Depth_+24 : Treatment_MAGICAL | -0.011 | -0.030 | 0.008 | 0.010 | -1.15 | 0.250 |

Random Effects :

| Group | Name | Variance | Std.Dev. |
|---|---|---|---|
| Mouse | (Intercept) | 0.025 | 0.158 |
| | Depth_000 | 0.000 | 0.008 |
| | Depth_+24 | 0.000 | 0.013 |
| Residual | | 0.159 | 0.398 |

Number of observations: 957264, groups: Mouse, 26

(**a**–**b**) Statistical results for the GLMMs with a Gamma error distribution and a log link function at CxLV (**a**) and HpCA1 (**b**). Wald-type 95 % CI was estimated for the coefficient of fixed effects by using the confint function.

**Supplementary Movie 1. *In vivo* two-photon imaging at mouse cortex between 450 and 900 µm depth.** The image stacks were captured with 3-µm z-step size by using two-photon microscopy with LP 66 mW and HV 30 at 1 fps. scale bar 100 µm. CxLV, cortical layer V; CxLVI, cortical layer VI; WM, white matter.
